# Multidimensional face representation in deep convolutional neural network reveals the mechanism underlying AI racism

**DOI:** 10.1101/2020.10.20.347898

**Authors:** Jinhua Tian, Hailun Xie, Siyuan Hu, Jia Liu

## Abstract

The increasingly popular application of AI runs the risks of amplifying social bias, such as classifying non-white faces to animals. Recent research has attributed the bias largely to data for training. However, the underlying mechanism is little known, and therefore strategies to rectify the bias are unresolved. Here we examined a typical deep convolutional neural network (DCNN), VGG-Face, which was trained with a face dataset consisting of more white faces than black and Asian faces. The transfer learning result showed significantly better performance in identifying white faces, just like the well-known social bias in human, the other-race effect (ORE). To test whether the effect resulted from the imbalance of face images, we retrained the VGG-Face with a dataset containing more Asian faces, and found a reverse ORE that the newly-trained VGG-Face preferred Asian faces over white faces in identification accuracy. In addition, when the number of Asian faces and white faces were matched in the dataset, the DCNN did not show any bias. To further examine how imbalanced image input led to the ORE, we performed the representational similarity analysis on VGG-Face’s activation. We found that when the dataset contained more white faces, the representation of white faces was more distinct, indexed by smaller ingroup similarity and larger representational Euclidean distance. That is, white faces were scattered more sparsely in the representational face space of the VGG-Face than the other faces. Importantly, the distinctiveness of faces was positively correlated with the identification accuracy, which explained the ORE observed in the VGG-Face. In sum, our study revealed the mechanism underlying the ORE in DCNNs, which provides a novel approach of study AI ethics. In addition, the face multidimensional representation theory discovered in human was found also applicable to DCNNs, advocating future studies to apply more cognitive theories to understand DCNN’s behavior.

## 1. INTRODUCTION

With enormous progress in artificial intelligence (AI), deep convolutional neural networks (DCNN) have shown extraordinary performance in computer vision, natural language processing, and complex strategy video games. However, the application of DCNNs runs the risk of amplifying social bias (Zou & Schiebinger, 2018). For example, a word embedding processing system may associates women with homemakers, or a face identification network may match non-white faces to inanimate objects, suggesting the existence of gender and race biases in DCNNs (Bolukbasi, Chang, Zou, Saligrama, & Kalai, 2016). Though the phenomenon of social bias has been widely recognized, the underlying mechanism of such bias is little known (Caliskan, Bryson, & Narayanan, 2017; Garg, Schiebinger, Jurafsky, & Zou, 2018). Here we asked how biased behaviors were generated in DCNNs.

Insight on human biases may help understanding DCNNs’ biases. A classical race bias, the other race effect (ORE) (Malpass, Roy, Kravitz, & Jerome, 1969; Valentine, 1991), shows that people are better at identifying faces of their own race than those of other races (Meissner & Brigham, 2001). One influential theory, the face multidimensional representation (MDS) theory, proposes that the ORE comes from the difference in representing faces in a multidimensional space, or simply ‘face space’ (O’Toole, Castillo, Parde, Hill, & Chellappa, 2018; Valentine, 1991; Valentine, Lewis, & Hills, 2016). According to this theory, face space is a Euclidean multidimensional space, with dimensions representing facial features. The distance between two faces in the space indexes their perceptual similarity. Under the frame of this theory, faces of one’s own race are scattered widely in the face space (i.e., high distinctiveness) and faces of other races are clustered in a smaller space (i.e., low distinctiveness) (Valentine, 1991; Valentine et al., 2016). Therefore, the higher distinctiveness in representation leads to better recognition of own-race faces than that of other-race faces. In this study, we examine whether the ORE in DCNNs, if observed, may be accounted for by a similar mechanism.

To address this question, current study chose a typical DCNN, VGG-Face (Figure 1A), which is widely used for face recognition (Parkhi, Vedaldi, & Zisserman, 2015). We first examined whether there was a similar ORE in the VGG-Face, and then explored its face representation space with the MDS theory. Specifically, we first manipulated the ratio of face images of different races to examine whether the ORE in the VGG-Face changed as a function of the frequencies of encountered races (Chiroro & Valentine, 1995). Furthermore, we examined whether frequent interaction of one race leads to sparser distribution (i.e., high distinctiveness) in VGG-Face’s representation space. Finally, we explored whether the difference in representation led to the ORE.

**FIGURE 1.**
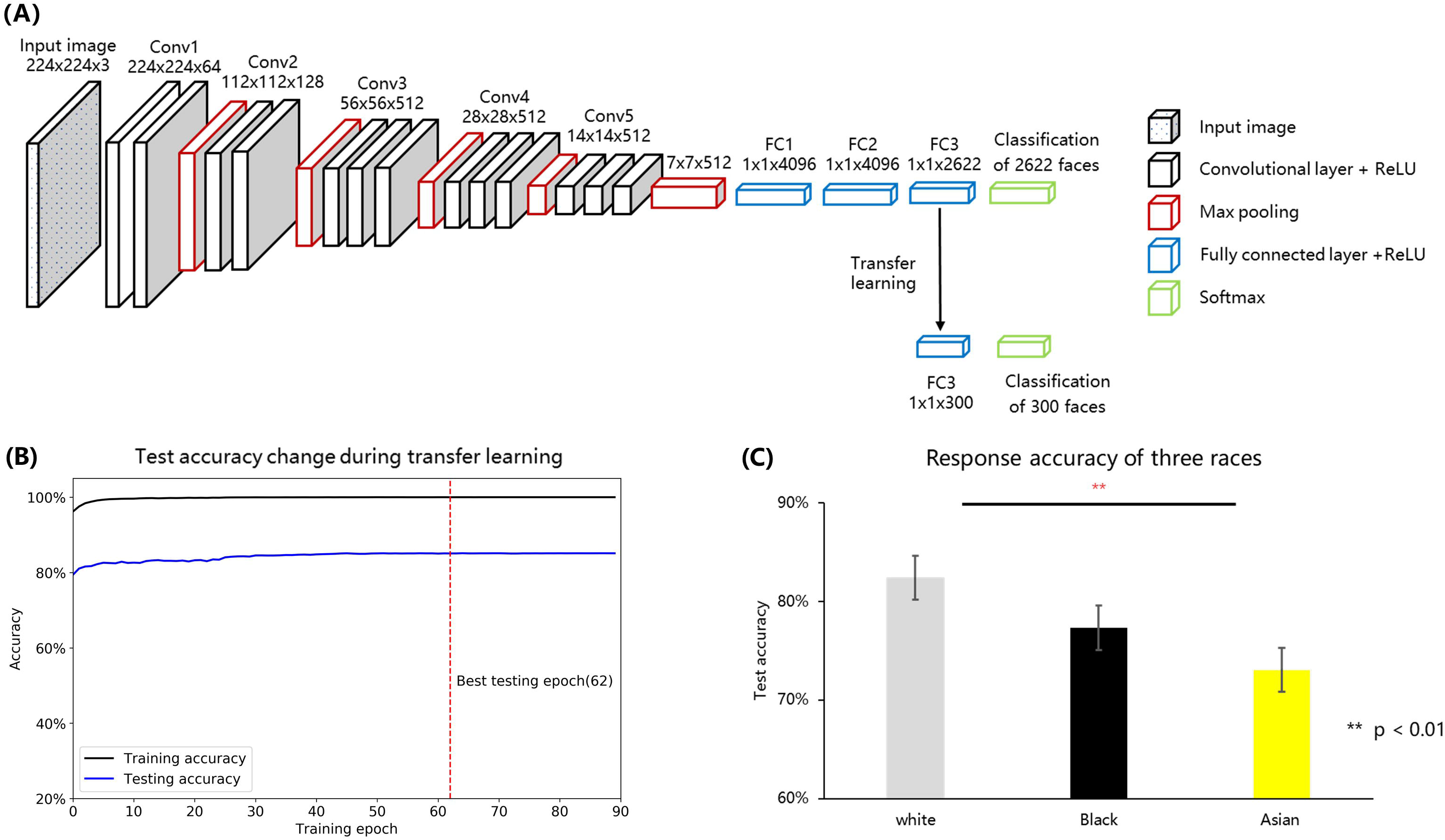
**(A)** Illustration of VGG-Face’s architecture used in this study. The model comprised five convolutional blocks (conv1-conv5) and three fully connected layers (FC1-FC3). **(B)** The change of test accuracy during VGG-Face transfer learning. The x axis represents for training accuracy, and the y axis represents for training epochs. The black and blue line represent for training and validation accuracy changes during model training separately. **(C)** Identification accuracy of the VGG-Face on white, black, and Asian faces.

## 2. MATERIALS AND METHODS

### 2.1 Convolutional neural network model

In this study, a well-known deep neural network, VGG-Face (available in http://www.robots.ox.ac.uk/~albanie/pytorch-models.html) was used for model testing, model retraining, and model activation extraction (Parkhi et al., 2015). Illustrations of VGG-Face architecture is shown in Figure 1A. This framework consists of 5 groups of convolutional layers and 3 fully connected layers, 16 layers in total. Each convolutional layer comprises some convolution operators, followed by non-linear rectification layer such as ReLU and max pooling. The input images (for example, 3 × 224 × 224 pixel color image) are transferred into 2622 representational units, each corresponding to a unit of the last fully connected layer (FC3) and representing for a certain identity.

### 2.2 Face stimuli

The VGG-Face was originally trained for face identification task with the VGGFace dataset (2622 identities in total, and 2271 identities is downloadable).

To test the performance of the VGG-Face on 3 races, 300 different identities were selected from another face dataset -- VGGFace2 (Cao et al., 2018). Faces images that are present in both VGGFace and VGGFace2 datasets were all excluded (see http://www.robots.ox.ac.uk/~vgg/data/vgg_face2/meta/class_overlap_vgg1_2.txt for details). We classified the remaining 8250 identities into four groups, white (6995 identities), black (518 identities), Asian (345 identities), and other races (392 identities). Three hundred identities were randomly selected from the first three groups (100 identities for each race) and separated into in-house training (300 identities, each containing 100 images), validating (300 identities, each containing 50 images) and testing (300 identities, each containing 50 images) datasets. These three datasets contained the same identities but with different face exemplars. We trained the VGG-Face with the training dataset, and validated the model with the validating dataset, and finally used the testing dataset to measure the identification accuracy of three different races.

### 2.3 Transfer Learning

We trained the VGG-Face with new identities using transfer learning (Yosinski, Clune, Bengio, & Lipson, 2014), which is to train a pretrained network with another small set of related stimuli. Transfer learning was performed to the pretrained VGG-Face with the in-house training set. We replaced the last FC layer (the third fully connected layer, FC3, containing 2,662 units) of the VGG-Face with another fully connected layer containing 300 units (each represent for a unique face identity used in training and testing procedure). Then, we froze the parameters prior to the classification layer (FC3), and trained the FC3 using the training dataset. Detailed training parameters were from a previous study (Krizhevsky, 2014). All networks are trained for face identification using the cross-entropy loss function with stochastic gradient descent (SGD) optimizer (initial learning rate = 0.01, momentum = 0.9). Images were normalized to the same luminance (mean = [0.485, 0.456, 0.406], SD = [0.229, 0.224, 0.225]) and resize to the 3 × 224 × 224 pixels. Data argumentation used 15° random rotation and 50% chance of horizontal flip. All models were trained for 90 epochs, and the learning rate decayed 250^−1/3^ (≈ 0.159) after each 23 epochs (1/4 training epochs). To achieve best training accuracy and prevent from overfitting, we save the ‘best model’ during training, referring to the best validating accuracy. The training procedure is shown in Figure1B. After the transfer learning, this network (the best model) was tested using the testing dataset. The performance difference between the three races was analyzed using repeated-measures analysis of variance (ANOVA).

### 2.4 Model retraining

According to human contact theory, low interracial interaction is the main cause of the ORE. For a DCNN, biased training data may lead to biased performance. To examine this hypothesis, we further retrained the VGG-Face using two ‘biased’ face sets and one matched face set, and then tested whether these models show the face bias. The training face sets were composed of different amounts of Asian and white faces. The different composition of Asian and white faces simulates the “white biased,” “Asian biased,” and “unbiased” datasets.

Re*training materials* All images used for model retraining and validating were also selected from VGGFace2 datasets. We selected 404 Asian identities and 404 white identities for model training and testing. For the white-biased model, we randomly selected 304 white identities out of 404 identities for model training. For the Asian-biased model, we randomly selected 304 Asian identities out of 404 identities for model training. For unbiased model training, we selected 152 Asian and 152 white identities. The training datasets were further separated into training and validating sets. We selected 30 of each identity (15000 images in total) as the validating dataset, and the remaining faces (109450 images for Asian biased model, 103745 images for white biased model, and 105781 images for unbiased model) were used for model training. Two hundred other identities (100 identities for each race) were selected for transfer learning and testing.

*Retraining procedure* We used the same VGG-Face framework as the pretrained model. All networks were trained for face identification using the cross-entropy loss function with stochastic gradient descent (SGD) optimizer (initial learning rate = 0.01, momentum = 0.9). Images were normalized to the same luminance (mean = [0.485, 0.456, 0.406], SD = [0.229, 0.224, 0.225]), and resized to 3 × 224 × 224 pixels. Data argumentation used 15° random rotation and 50% chance of horizontal flip. All models were trained for 90 epochs, learning rate decayed 250^−1/3^ (≈ 0.159) after each 23 epochs (1/4 training epochs). To achieve best training accuracy and prevent from overfitting, we save the best model during training, referring to the best validating accuracy. The saved model was used for further model testing using testing dataset.

### 2.5 Face representation difference of three races in VGG

To explore the representation pattern of different races in the VGG-Face, we further analyzed the face representation difference. It has been suggested that activation responses of the layer prior to the final classification layer (the second fully connected layer, FC2) is the typical representation of each face in DCNNs (O’Toole et al., 2018). Thus, we extracted the activation responses in the FC2 for all the testing faces using an in-house python package DNNBrain (Chen et al., 2020) with the PyTorch framework (Paszke et al., 2019).

To describe the distinctness of each race groups, we used three measurements to describe the distribution of face space. First, we applied the representation similarity analysis to get the representational dissimilarity correlation matrix (RDM) of three race faces with FC2 activation. To further explore the representation difference of three races, we used the ingroup similarity to describe the representation variance within a race group. Ingroup similarity was calculated as the averaged Pearson correlation of a certain identity with other identities of the same race. Specifically, a face with larger ingroup similarity indicates smaller representation distinctiveness. That is, the larger the distinctiveness is, the better the performance in discriminating identities is.

Next, we used the FC2 activation to construct the face space that describes the distribution of different faces. Valentine and Endo (1992) assume the face space to be an n-dimensional space, a face is represented as a point localized in the space. The axes of the space represent for dimensions to discriminate faces. According to this hypothesis, we used average activation of all faces as the possible center coordinates of this face space. Thus, we compute Euclidean distance of the averaged activation from each face to all averaged face activations as a measurement of face distinctiveness. A face with larger Euclidean distance indicates larger representation distinctiveness. The activation differences in the three races were also analyzed using one-way ANOVA.

### 2.6 Face representation visualization

For a better visualization of the representation of the face space, we used the t-SNE (t-distributed stochastic neighbor embedding, t-SNE) method to reduce face representation dimensions and visualize the activation distribution. The t-SNE starts by converting the high-dimensional Euclidean distances between datapoints into conditional probabilities that represent similarities (Maaten & Hinton, 2008). We used the t-SNE to squeeze the activation vectors (2622 units) of each face’s activation into two dimensions and plot these conditional probabilities on a two-dimensional coordinate for visualization. The t-SNE was performed using default parameters (learning rate = 200, iteration = 1000).

### 2.7 Correlation between face representation and identification performance

To explore whether VGG-Face’s activation and its performance was correlated, we computed the Spearman correlation as well as the Pearson correlation between the ingroup similarity and Euclidean distance with face identification accuracy of the VGG-Face.

## 3. RESULTS

First, we used transfer learning to examine the race bias in the VGG-Face. The averaged accuracy of all identities was 77.6%, significantly higher than the stochastic probability (0.33%), indicating the success of transfer learning. A one-way ANOVA showed a significant main effect of race (*F*_2,297_ = 8.762, *p* < 0.001, 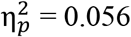), with white faces identified significantly better than Asian (*p* < 0.001, *d*^*’*^ = 0.545) and marginal significantly better than black (77.32% ± 14.32%, *p* = 0.071, *d*^*’*^ = 0.353) faces (Figure 1C). No significant difference was found in accuracy between black and Asian faces (*p* = 0.176).

To verify face selection bias in VGG network training, we classified the available VGGFace dataset into 4 groups, that is, white (1984 identities, 87.2%), black (211 identities, 9.7%), Asian (52 identities, 2.3%), and other races (brown or mixed race, 24 identities, 1.1%). For faces in the dataset were overwhelmingly from white, the better identification accuracy for white faces suggested that the ORE also existed in the VGG-Face.

A direct test on whether the ORE observed in the VGG-Face resulted from the imbalance of races present in the dataset was to manipulate the ratio of the number of each race faces. To do this, we retrained the VGG network using white-biased (White versus Asian: 100% versus 0%), Asian biased (0% versus 100%), and unbiased (50% versus 50%) datasets, respectively. As shown in Figure 2, three DCNNs showed different patterns of ORE. For the DCNN trained with the white-biased dataset, white faces were identified significantly better than Asian faces (t_198_ = 3.934, *p* < 0.001, *d*^*’*^ = 0.562). In contrast, in the Asian-biased DCNN, Asian faces were identified better than white faces (t_198_ = 2.693, *p* = 0.008, *d*^*’*^ = 0.381). Finally, no ORE was found in the unbiased DCNN (t_198_ = 1.135, *p* = 0.258, *d*^*’*^ = 0.161). Taken together, the ORE observed in the VGG-Face resulted from unbalanced experiences with different number of faces per race during model training.

**FIGURE 2.**
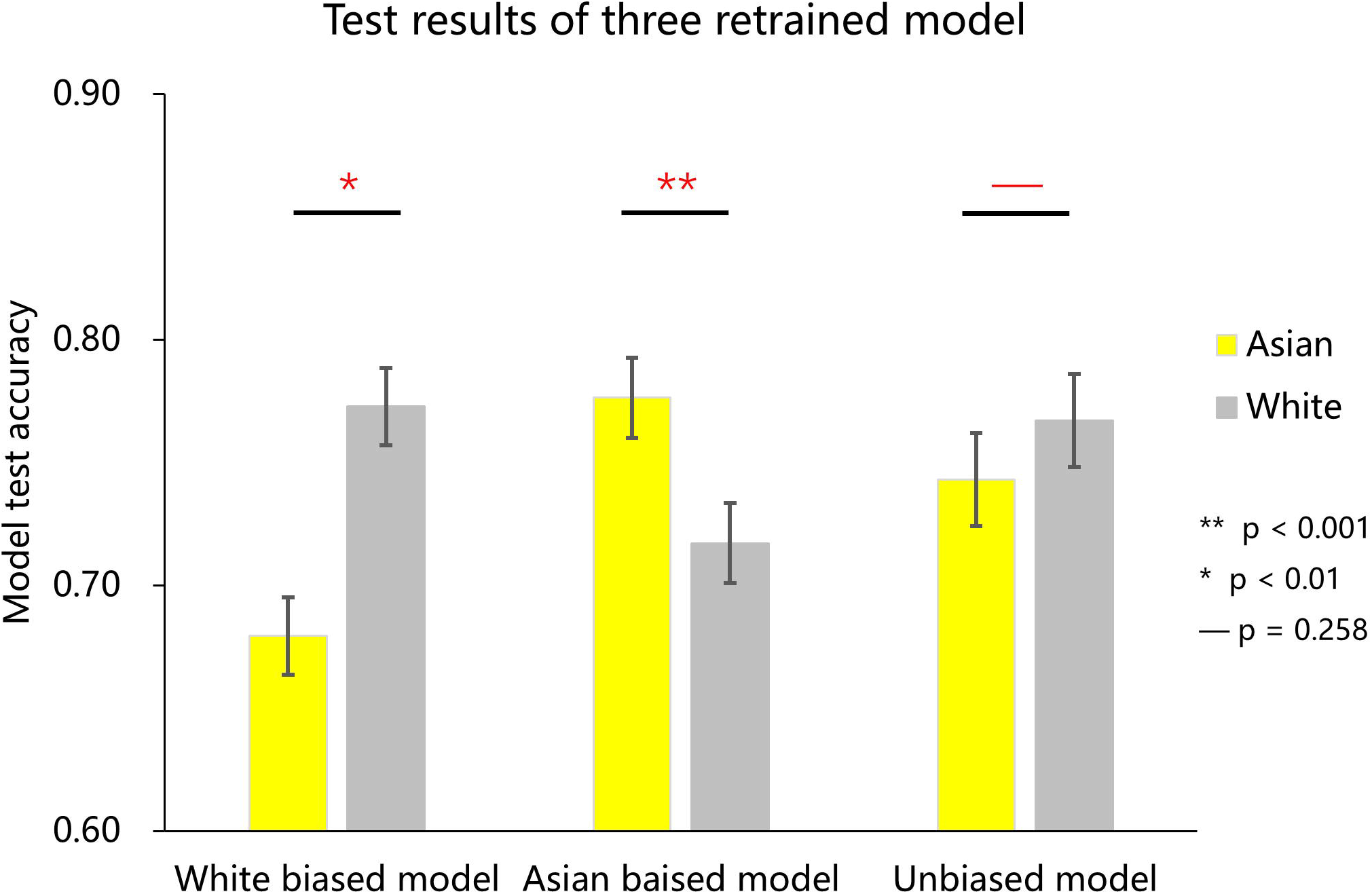
Identification performance of three retrained VGG networks, that is, White biased model, Asian biased model, and unbiased model.

How did unbalanced experiences shape the internal representation for faces in the VGG-Face? To address this question, we calculated correlations between the representations of faces, which were indexed by the activations in FC2 layer, and then constructed the correlation matrix consisting of Asian, white, and black faces (Figure 3A). A direct observation of Figure 3A revealed that faces of each race were grouped into one cluster; that is, the representations for faces were more similar within a race than between races, suggesting that faces from the same race were grouped together in the multidimensional space. Importantly, the representational similarity of white faces was smallest as compared to that of Asian (*p* < 0.001, *d*^*’*^ = 1.29) and black (*p* < 0.001, *d*^*’*^ = 2.077) faces, and that of Asian faces was smaller than that of black faces (*p* < 0.001, *d*^*’*^ = 0.4) (Figure 3B). That is, the representations for white faces were the sparsest in the face space. To quantify the sparseness of the representation, we calculated the Euclidean distance of the representation of individual faces to the averaged representation of all faces. As shown in Figure 3C, the representation of white faces was localized farther from the averaged representation than that of Asian (*p* = 0.008, *d*^*’*^ = 0.386) and black (*p* < 0.001, *d*^*’*^ = 1.286) faces, and that of Asian faces was farther than that of black faces (*p* < 0.001, *d*^*’*^ = 0.773). Activation of faces in the last fully connected layer (FC3) was also extracted and analyzed, which showed similar representational pattern as FC2 (detailed information is provided in the supplementary materials).

**FIGURE 3.**
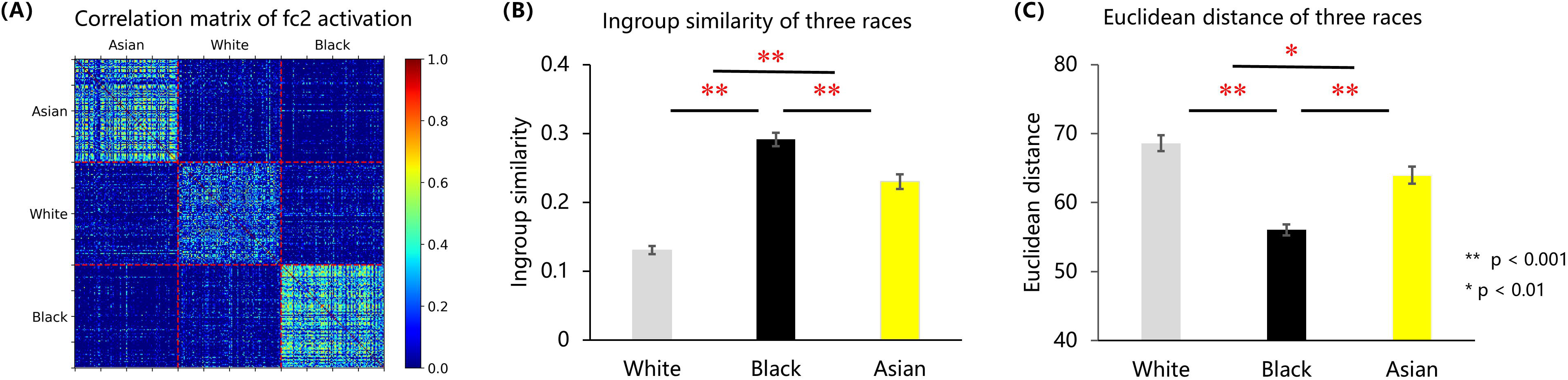
**(A)** VGG-Face FC2 activation correlation matrix of Asian, white, black faces. (**B)** Face distinctiveness of white, black, and Asian faces measured using ingroup similarity. **(C)** Face distinctiveness of white, black, and Asian faces measured using face Euclidean distance.

To visualize how race faces were represented in the face space, we used t-SNE to reduce multiple dimensions to 2 dimensions. As shown in Figure 4A, representations for each race was grouped into one cluster; however, the clusters for Asian and black faces were denser, whereas white faces were distributed sparser in the face space.

**FIGURE 4.**
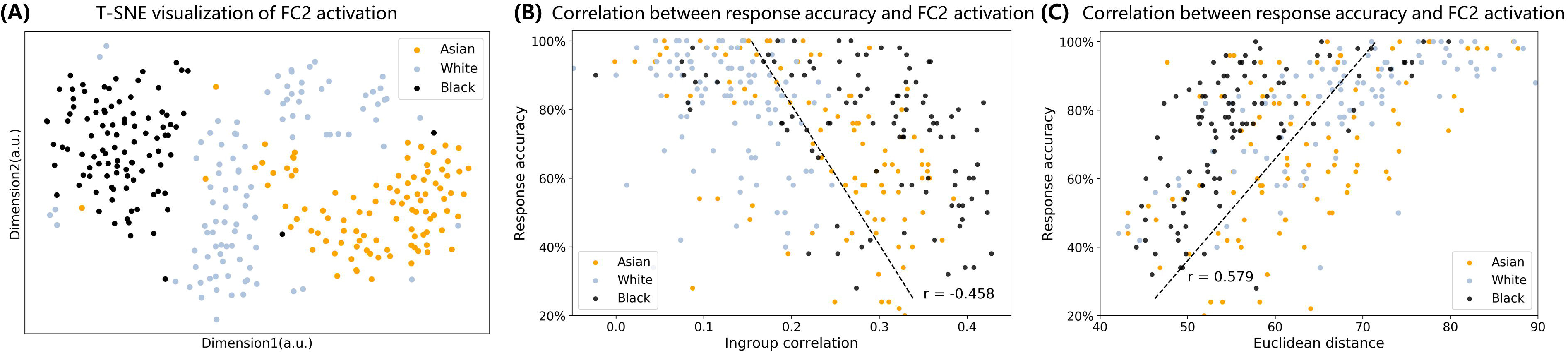
**(A)** T-SNE visualization of FC2 activation of Asian, white, black faces. **(B)** Correlation between ingroup similarity and face identification accuracy. **(C)** correlation between face Euclidean distance to averaged face activation and face identification accuracy.

Finally, we asked whether the difference in sparseness of the representation related to the ORE observed in VGG-Face. As shown in Figure 4B, the correlation analysis showed a significant negative correlation between ingroup similarity and face identification accuracy (coefficient Pearson’s R = −0.458, *p* < 0.001, coefficient Spearman correlation R = −0.499, *p* < 0.001). As shown in Figure 4C, the correlation analysis showed a significant positive correlation between Euclidean distance and face identification accuracy (coefficient Pearson’s R = 0.579, *p* < 0.001, coefficient Spearman correlation R = 0.621, *p* < 0.001). That is, if a face was represented farther from the averaged representation, it was more accurately identified by the VGG-Face. For the VGG-Face trained by a dataset dominated by white faces, white faces on average had the largest representational distance, and they were the most likely to be identified correctly, which therefore resulted in the ORE.

## 4. DISCUSSION

In this study, we examined the ORE in VGG-Face. By manipulating the ratio of faces of different races in the training dataset, the result demonstrated that unbalanced datasets led to the ORE in VGG-Face, in line with findings in humans that visual experiences affects the identification accuracy of race faces (Chiroro & Valentine, 1995; Meissner & Brigham, 2001). Importantly, the representation similarity analysis revealed that if white faces dominated the dataset, they scattered more sparsely in the multidimensional representational space of faces in VGG-Face, and thus resulted in better behavioral performance. In sum, with the MDS theory in human, we provided a novel approach to understand biases in DCNNs.

The AI ethic problem has attracted broad attention in the field of AI (Zemel, Wu, Swersky, Pitassi, & Dwork, 2013; Zou & Schiebinger, 2018). However, the mechanism underlying AI biases is poorly understood. Our study confirmed that bias may derived from unbalanced training dataset. It’s coincident with the contact theory (Chiroro & Valentine, 1995) in humans, in which high-contact faces are recognized more accurately than that of low-contact ones. Previous studies in humans suggest that high ingroup interaction leads to sparser representation (high distinctiveness) of ingroup faces in face space, whereas low interaction leads to denser representation (low distinctiveness) of outgroup faces (Valentine, 1991; Valentine et al., 2016). In current study, we also found that in the representational space of VGG-Face, the “own-race” face (i.e., white faces) showed larger distinctiveness than that of “other-race” faces (i.e., Asian, and black faces). Furthermore, the distinctiveness was indexed by the representational similarity of faces, which may serve as a more sensitive index than the ratio of faces in the unbalanced dataset. Therefore, before the formal training, an examination of representational similarity in MDS with a portion of the training dataset may provide an estimate of the skewness of the datasets and the biased performance under current task demands.

Therefore, a more effective way of controlling AI biases may come from new algorithms that can modulate internal representations of DCNNs. Currently, the main effort has been focusing on the construction of balanced datasets and the approaches of training DCNNs, and guidelines have been advised (Gebru et al., 2018; Mitchell et al., 2019). However, it is laborious to balance datasets not only in terms of data collection, but also in terms of task demands. It might be more efficient if a revised back-propagation algorithm could both minimize errors between outputs and goals, and rectify differences in distinctiveness of the representation of interests. For example, in the field of natural language processing, Beutel (Beutel, Chen, Zhao, & Chi, 2017) and Zhang (Zhang, Lemoine, & Mitchell, 2018) propose a multi-task adversarial learning method to manipulate the biased representational subspace and thus mitigate gender bias of model performance. They build a multi-head DCNN where one head is for target classification and another head is for removing information about unfair attribute learned from the data. Similarly, in the field of computer vision, further study could also explore the way to manipulate the face representational space to reduce social bias in DCNNs.

In conclusion, our study used a well-known phenomenon, the ORE, to investigate the mechanism inside DCNNs that leads to biased performance. In addition, we found a human-like multidimensional face representation in DCNN, suggesting that paradigms and theories discovered in human studies may also be helpful to unveil the black box of DCNNs. There are many other types of biases in AI such as gender bias and age bias, and therefore our study invites broad investigation on these ethic problems in AI.

## DATA AVAILABILITY STATEMENT

All face image materials and model training codes used in this article are provided on git-hub (https://github.com/JinhuaTian/DCNN_other_race_effect).

## ACKNOWLEDGEMENTS

This study was funded by the National Natural Science Foundation of China (31861143039) and the National Basic Research Program of China (2018YFC0810602).

