## supplementary materials for "Multidimensional face representation in deep convolutional neural network reveals the mechanism underlying AI racism"

### *Supplementary Material*

#### 1.1 Face-race representation difference of three races in FC3

As for fc3, One-way ANOVAs results showed significant main race effect in ingroup similarity ($F_{2,297}$ = 121.327, $p$ < 0.001, $\eta_p^2$ = 0.450). As shown in Figure 1A, pairwise analysis (with Bonferroni correction) indicated that white faces showed significantly smaller ingroup similarity (larger distinctiveness) than Asian ($p$ < 0.001, $d'$ = 0.863) and black ($p$ < 0.001, $d'$ = 2.590) faces, and Asian faces showed smaller ingroup similarity than black faces ($p$ < 0.001, $d'$ = 1.222). This result indicated that white faces showed less ingroup representational similarity than both Asian and black faces in VGG network.

Moreover, one-way ANOVAs results also showed a significant main race effect in both Euclidean distance ($F_{2,297}$ = 36.994, $\eta_p^2$ = 0.199, $p$ < 0.001). As shown in Figure 1B, pairwise comparison (with Bonferroni correction) showed that white faces were localized farther than Asian ($p$ < 0.001, $d'$ = 0.528) and black ($p$ < 0.001, $d'$ = 1.222) faces, and Asian faces were localized farther than black faces ($p$ < 0.001, $d'$ = 0.763).

These results indicated that fc3 showed similar representational pattern as fc2. That is, white faces showed larger distinctiveness (smaller ingroup similarity and larger Euclidean distance) than Asian and black faces.

#### 1.2 Correlation between face representation and identification performance in FC3

For FC3 activation, As shown in Figure 1C, correlation analysis showed a significant negative correlation (coefficient Pearson's R = -0.368, $p$ < 0.001, coefficient Spearman correlation R = -0.420, $p$ < 0.001) between ingroup similarity and face identification accuracy. As shown in Figure 1D, correlation analysis also showed a significant positive correlation between Euclidean distance (coefficient Pearson's R = 0.451, $p$ < 0.001, coefficient Spearman correlation R = 0.478, $p$ < 0.001) and VGGFACE identification accuracy. FC3 results showed similar activation-performance correlation-ship as FC2, which indicated that a face with higher identification accuracy (larger face distinctiveness) showed smaller ingroup representational similarity with other faces, and localized farther in the face space.

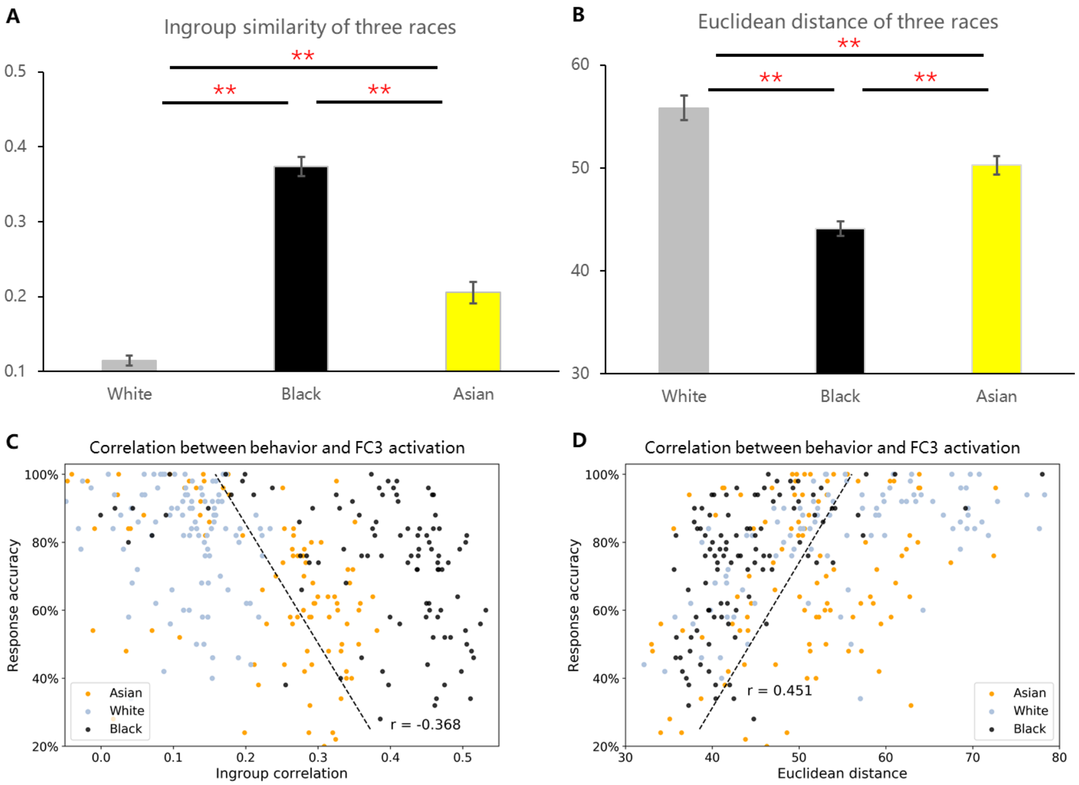


**FIGURE 1. | (A)** Face distinctiveness of Asian, white, and black faces measured using ingroup similarity. **(B)** Face distinctiveness of Asian, white, and black faces measured using face Euclidean distance to all averaged face activation. **(C)** correlation between face distinctiveness and VGG face identification accuracy in FC3. **(D)** correlation between Face Euclidean distance and VGG face identification accuracy in FC3.

#### 1.3 Model retraining details

Train procedure of white biased model, Asian biased model, and unbiased model are shown in Figure 2A, 2B, and 2C. For white biased model, its best validating epoch is epoch 85 and the best validating accuracy is 0.989. For Asian biased model, its best validating epoch is epoch 67 and the best validating accuracy is 0.962. For unbiased model, its best epoch is epoch 61 and the best validating accuracy is 0.975.

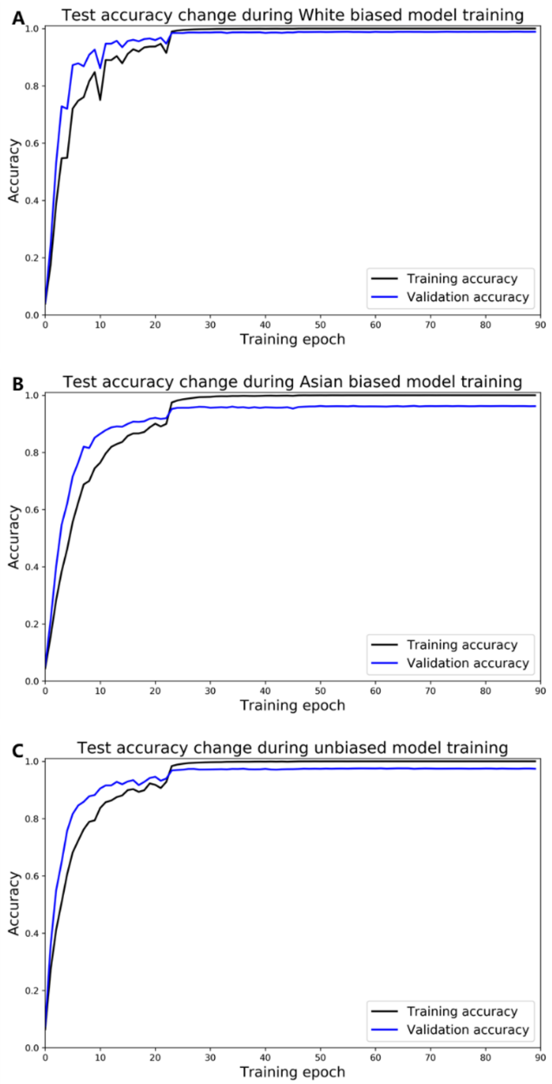


**FIGURE 2 | (A)**, **(B)**, **(C)**, training, and validating procedure during the model training of white biased model, Asian biased model, and unbiased model.
